# Functional segregation of cortical hand and speech areas by frequency detuning of an intrinsic motor rhythm

**DOI:** 10.1101/2024.11.11.622882

**Authors:** Ioanna Anastasopoulou, Douglas O Cheyne, Blake W Johnson

## Abstract

Many decades after Penfield’s (1937) classic depiction of the motor homunculus, it remains unclear how spatially contiguous and interconnected representations within human sensorimotor cortex might separate their activities to achieve the directed and precise control of distinct body regions evident in activities as different as typing and speaking. One long-standing but relatively neglected explanation draws from models of simple physical systems (like swinging pendulums) to posit that small differences in the oscillatory properties of neuronal populations (termed “frequency detuning”) can result in highly effective segregation of their activities and outputs. We tested this hypothesis by comparing the peak frequencies of beta-band (13-30 Hz) motor rhythms measured in a magnetoencephalographic neuroimaging study of finger and speech movements in a group of healthy adults and a group of typically developing children. Our results confirm a peak frequency task difference of about 1.5 Hz in the beta motor rhythms of both left and right hemispheres in adults. A comparable task difference was obtained in children for the left but not for the right hemisphere. These results provide novel support for the role of frequency detuning in the functional organisation of the brain and suggest that this mechanism should play a more prominent role in current models of bodily representations and their development within the sensorimotor cortex.

**NEW & NOTEWORTHY:** This work is the first evidence for frequency detuning of an intrinsic motor rhythm in spatially contiguous regions of primary motor cortex associated with hand and speech movements, supporting a mechanism of functional segregation that has been largely overlooked by current models of motor cortex organization. Our results from children further provide the first evidence for frequency detuning in the developing brain, indicating an early-developing mechanism with later refinements of hemispheric control.

## INTRODUCTION

Our understanding of the representational organization of human primary motor cortex has been guided for much of the last century by Penfield and Boldrey’s (1) classic diagram of a motor homunculus, a simple and conceptually appealing solution for the problem of organizing motor control of the body onto a flattened cortical surface. However it has long been realized that the Penfield homunculus is oversimplified and incomplete, and at the present time this model is undergoing vigorous revision to incorporate new findings that sharply depart from an elementary mapping of the body onto the cortical surface (2–6); and explanations of high dimensional motor behaviors that cannot be accommodated within a two-dimensional framework (7). This ongoing reconceptualization is being driven by new insights into cortical function and organization based on methodological advances in structural and functional brain mapping and brain connectivity techniques (8, 9), and new data from invasive and noninvasive recordings of neuronal function, and invasive and noninvasive brain stimulation techniques (10).

One core concept of functional brain organization has thus far played little role in these recent investigations of primary motor cortex (M1) organization and remains largely absent from modern models of M1 representation and function (11). It is now well-established that neurons exhibit oscillatory behaviors when engaged in cognitive, sensory and motor processes, and that these oscillations exhibit synchronization across neuronal populations at characteristic frequencies within both local circuits and larger brain networks (12). Neural network models posit that synchronous neural oscillations are a critical mechanism for establishing neural communication and cooperation, while in contrast, the existence of a frequency difference (termed “frequency detuning”) between oscillators acts against their synchronization by causing their phase relationship to change continuously (phase precession).

The existence of frequency differences (“frequency detuning”) among different body part representations points to an important organizing principle of sensorimotor cortex. Indeed, the extant literature provides good albeit very limited evidence for frequency detuning among individual motor representations within both M1 cortical and subcortical motor structures. In their EEG study Neuper and Pfurtscheller (13) reported have reported distinct peak resonance frequencies within the beta-band motor rhythm (13-30 Hz) for upper versus lower limb sensorimotor representations, with significantly higher peak beta frequencies (> 20 Hz) at recording locations over the lower limb representation compared to locations over the upper representation (< 20 Hz). A comparable result has been obtained for beta band upper versus lower limb representations in the subthalamic nucleus (14).

Apart from the electrophysiological studies described above there currently exists little empirical evidence for frequency detuning among different body part representations in M1 or other brain motor networks. In the present study we addressed this gap with a reanalysis of manual (button press) and speech elicited magnetoencephalographic data collected from healthy adults and typically developing children in studies previous described in (15, 16). Our analyses focused on deriving the peak beta frequencies for spatially adjacent regions of hand and speech sensorimotor cortices respectively.

## MATERIALS AND METHODS

Experimental methods are described in detail in (16) and summarised in brief below.

### Participants

Analyses were performed on data obtained from 10 adult participants (4F; mean age 32.5, range 19.7–61.8) and 19 typically developing (TD) children (8 females, mean age =11.0 years; SD = 2.5, range 7.5 – 16.7) and previously described in (16). All participants were right-handed as assessed by a short version of the Edinburgh handedness questionnaire (17). All procedures were approved by the Macquarie University Human Subjects Research Ethics Committee.

### MEG recordings

Neuromagnetic brain activity was recorded with a KIT-Macquarie MEG160 (Model PQ1160R-N2, KIT, Kanazawa, Japan) whole-head MEG system consisting of 160 first-order axial gradiometers with a 50-mm baseline (18, 19). MEG data were acquired with analogue filter settings of 0.3 Hz high-pass, 200 Hz low-pass, 1000 Hz sampling rate and 16-bit quantization. Measurements were carried out with participants in supine position in a magnetically shielded room (Fujihara Co. Ltd., Tokyo, Japan). For adult participants, MEG data were co-registered with individual T1-weighted anatomical magnetic resonance images (MRIs) acquired in a separate scanning session. Individual structural MRI scans were not available for the child participants in this study so a surrogate MRI approach was used which warps a template brain to each subject’s digitized head shape (20).

### Audio speech recordings

Time-aligned audio speech recordings were recorded in an auxiliary channel of the MEG setup with the same sample rate (1000 Hz) as the MEG recordings and were used to identify speech onset/offset events for use in MEG source reconstruction analyses.

### Experimental tasks

Participants were required to perform a manual (button press) task and a speech production task. For the manual task participants performed a button press task on a fibre optic response pad (Current Designs, Philadelphia) with the index finger of the dominant (right) hand at a self-paced rate of about 1 per 2 s for 180 s, resulting in a total of about 90 button press trials. For the speech task, participants performed a *reiterated nonword speech task* which limits requirements for semantic, syntactic and attentional processing (21) and emphasises motoric sequencing operations (22). Participants were required to produce utterances of the disyllabic V1CV2 sequences /ipa/ and /api/ at a constant rate during the course of a single exhalation of a deep breath intake. Each participant generated ten “trial sets” (23) for each speech condition, with each trial set lasting approximately 12 seconds. Each 12 sec trial set resulted in about 10 separate nonword utterances at the normal rate and about 15 at the faster rate. An inter-trial set rest interval of 4 secs was terminated by instructions (1 sec) for fixation and breath intake for the next trial set. Participants were instructed to minimize head movement and avoid blinking during speech trial sets.

### Data preprocessing and source reconstruction

Source reconstruction was performed using the scalar synthetic aperture magnetometry (SAM) beamformer implemented in the BrainWave MATLAB toolbox (24). For manual task data, MEG data were initially segmented into 1.5 s epochs comprising -0.5 sec to +1.0 sec with respect to button-press onset, resulting in about 90 button press trials. Epoched data sets were digitally filtered from 0.3-100 Hz and a 50 Hz notch filter. SAM source images were computed using a frequency range of 15-25 Hz (beta band), a sliding active window of 0.6 to 0.8 sec and a baseline window of -0.5 to -0.3 sec over 10 steps with a step size of 0.01 sec and pseudo-z beamformer normalisation (3 ft/sqrt (Hz) RMS noise).

For speech task data, a total of 40 speech trial set onsets were identified and marked from the speech channel of the MEG recordings. MEG data were segmented with an epoch of -10 sec to + 5 sec from the onset of each trial set. All epoched data were digitally filtered with a bandpass of 0.3-100 Hz and a 50 Hz notch filter. SAM source images were computed using a frequency range of 15-25 Hz, a sliding active window of 0.6 to 0.8 sec and a baseline window of -0.5 to - 0.3 sec over 10 steps with a step size of 0.01 sec and pseudo-z beamformer normalisation (3 ft/sqrt (Hz) RMS noise). The SAM noise-normalized (pseudo-t) beamformer images were computed using a frequency range of 15-25 Hz (center of the beta frequency band), a sliding active window of 0 to 1.0 sec (first second of current speech trial set) and a baseline window of -5 to -3 sec (first two seconds of inter trial set rest period) over 10 steps with a step size of 0.2 sec and pseudo-z beamformer normalisation (3 ft/sqrt (Hz) RMS noise).

Statistical analysis of group beamformer images was performed with cluster-based permutation testing (alpha = 0.05, 512-2048 permutations, omnibus correction for multiple comparisons). Detailed results of the beamformer source reconstructions are described in (15, 16). For visualisation purposes the results are depicted schematically in Figure 1, showing brain anatomical regions associated with significant left hemispheric beamformer clusters elicited by the manual and speech tasks respectively. For both adults and children, significant manual task clusters were centered on the hand knob of the precentral gyrus, an established anatomical landmark of the hand area of primary motor cortex (25) of the left cerebral hemisphere (contralateral to the right handed button press used in this task). All participants also showed smaller magnitude (and statistically non-significant at the group level) clusters in the homologous precentral gyrus region in the right hemisphere. For the speech task, the adult group showed a single significant beamformer cluster in the left hemisphere, centred at anatomical coordinates immediately inferior to the hand knob and encompassing the anatomical location of the medial frontal gyrus (MFG) and a region of the immediately adjacent precentral gyrus (middle precentral gyrus). In contrast, the child group showed bilateralized speech responses, with a significant cluster in each hemisphere.

**Figure 1.**
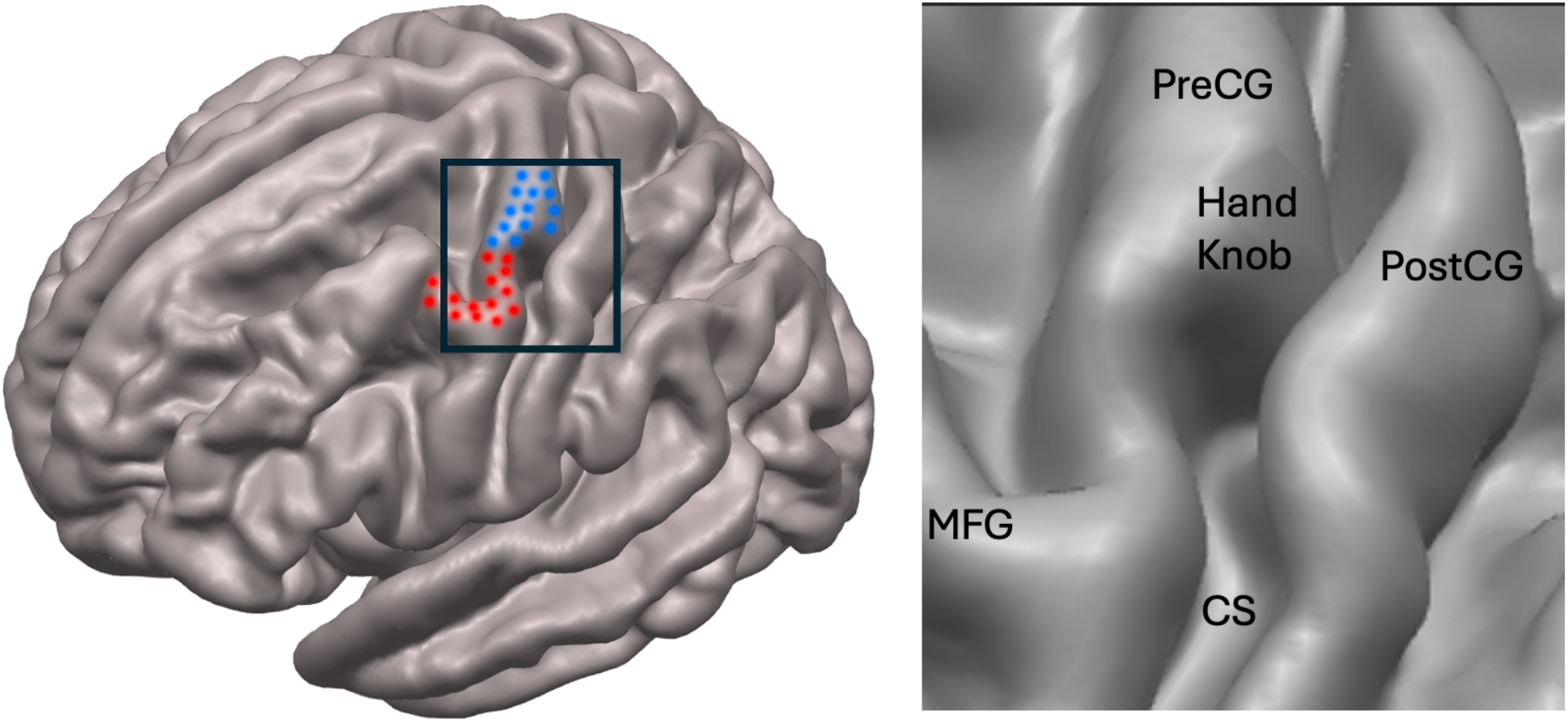
Manual and speech motor regions (left cerebral hemisphere). **Left**. Blue and red stippling indicate anatomical locations of SAM beamformer clusters associated with manual and speech tasks respectively (for left hemisphere of adult group). **Right**. Enlargement of bounded region of left panel shows anatomical landmarks PreCG = precentral gyrus, PostCG = postcentral gyrus, CS = central sulcus, MFG = medial frontal gyrus. Adapted from beamformer cluster maps of Figure 3 in (15).

### Peak frequency analysis

For the manual task data, voxel locations at the centre of group mean clusters were used to generate virtual sensor time frequency spectrograms for each trial with a time range of -0.4 to +0.9 sec from the button press onset and a frequency range of 1-100 Hz. For speech task data time frequency plots were generated with a time range of -9 to +4 sec from the trial set onset and a frequency range of 1-100 Hz. Time-frequency data for each individual was frequency averaged over the beta frequency band of 13-30 Hz. Based on the group-averaged beta-band power time series (Figure 2), analysis time-epochs were selected as: for the manual task, -0.2 to 0 sec (event-related desynchronization, ERD) and 0.6 to 0.8 sec (event-related synchronization, ERS) with respect to button press onset; and for the speech task, -5 to -3 sec (ERS) and -1 to 1 sec (ERD), with respect to speech trial set onset.

**Figure 2.**
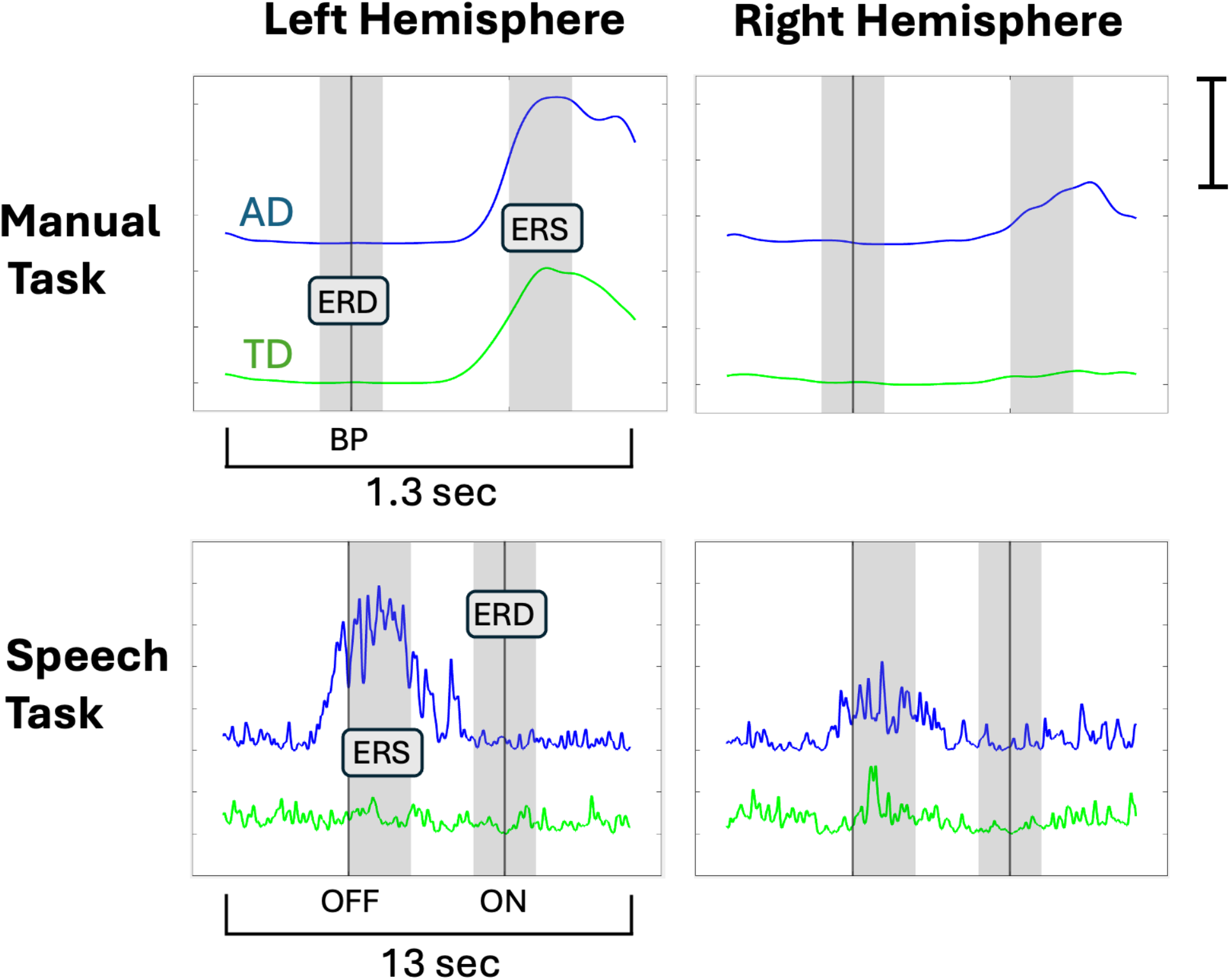
Time course of manual and speech beta-band responses in adults and children. Vertical black line indicates button press (BP) onset for manual task, speech offset (OFF) and onset (ON) for speech task. Shaded regions show time epochs used for computations of peak frequencies. ERD = event-related desynchronization epoch, ERS = event-related synchronization epoch, LH = left hemisphere, RH = right hemisphere. Scale bar indicates 25% power change from baseline. Adapted from data of Figure 4 in (16).

Peak beta frequency was estimated from the 13-30 Hz frequency region of trial-by-trial spectrograms for each individual and epoch. A bootstrap procedure with 2,000 iterations was used to obtain a minimum (ERD epochs) or maximum (ERS epochs) frequency value computed over the full epoch for each sample and to build a distribution whose mean was designated as the peak frequency. Each iteration of the bootstrapping procedure was a unique sample (without replacement) of M samples from N trials, with M = N/2 (26); see also (27) for a similar peak estimation approach).

### Statistical analyses

For each participant group, a 2×2×2 repeated measures ANOVA was computed with the factors of task (manual, speech), hemisphere (left, right) and epoch (ERS, ERD). Degrees of freedom were adjusted with the Greenhouse-Geisser correction where appropriate. Between group comparisons were performed separately for manual and speech tasks usings nonparametric Kruskal-Wallis one-way ANOVA. For each task (button press, speech), hemisphere (left and right) a linear regression model (MatLab *fitlm* function) was used to test for linear effects of age within the child group.

## RESULTS

Results of the peak frequency analyses are summarized in Figure 3.

**Figure 3.**
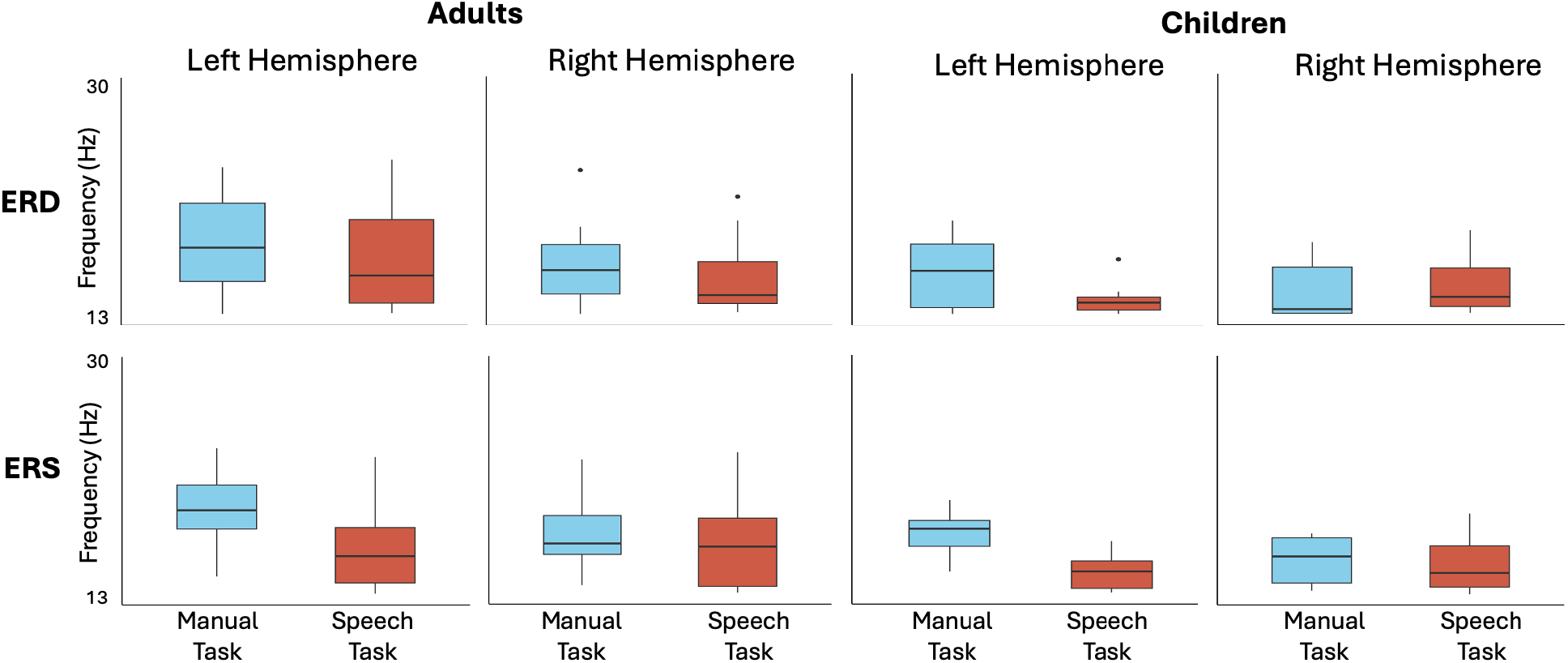
Estimated peak frequencies of beta-band response to manual and speech movements. Horizontal lines indicate median frequency, boxes indicate interquartile range, whiskers indicate range. For adults, ANOVA showed significant main effects of task and hemisphere. For children, ANOVA showed a significant task x hemisphere interaction.

Data for the adult group showed higher mean peak beta band frequency for the manual task relative to the speech task, for both left and right hemispheres and both ERD and ERS epochs. Overall mean peak frequencies (collapsing across hemisphere and epoch) were 17.883 Hz for the manual task and 16.583 Hz for the speech task, for a mean task difference of 1.300 Hz. ANOVA confirmed a significant main effect of task (F(1,9) = 8.692, p = .016, η^2^_p_ = .491). ANOVA also showed a significant main effect of hemisphere (F(1,9) = 9.779, p = .012, η^2^_p_ = .521). Visual comparison of the left and right hemisphere data indicates higher peak beta frequencies in the left hemisphere (*M* = 17.814 Hz) relative to the right hemisphere (*M* = 16.653 Hz), for a mean hemispheric difference of 1.161 Hz.

The three-way ANOVA showed no significant main effect of epoch (p = .107) and no significant interactions among the hemisphere, task, and epoch factors.

For the child group, Figure 3 shows that a Manual-Speech task difference is also apparent in the child group for the left hemisphere (manual *M* = 16.609 Hz, speech *M* = 14.292 Hz, difference = 2.316 Hz) but not for the right hemisphere (manual *M* = 15.125 Hz, speech *M* = 15.116 Hz, difference = 0.010 Hz). ANOVA confirmed a significant hemisphere x task interaction (F(1,18) = 23.774, p < .001, η^2^_p_ = .569). Post hoc contrasts confirmed a significant Task effect for the left hemisphere (p < .001) but not for the right hemisphere (p = .968).

The three way ANOVA also showed a significant main effect of epoch (F(1,18) = 8.330, p = .010, η^2^_p_ = .316), indicating that ERS peak frequency was significantly greater than ERD frequency (ERD M = 14.838, ERS M = 15.733, difference = 0.900 Hz), as well as a significant main effect of Task (F(1,18) = 31.939, p < .001, η^2^_p_ = .640). There was no significant main effect of hemisphere (p = .210) and no other significant interactions between the factors.

### Effects of age in child group

Figure 4 plots estimated peak frequency against age for individual children. No significant linear effects of age were obtained for obtained for either frequency band or hemisphere. Task effects are clearly evident for beta band peak frequencies in the left hemisphere but not in the right hemisphere.

**Figure 4.**
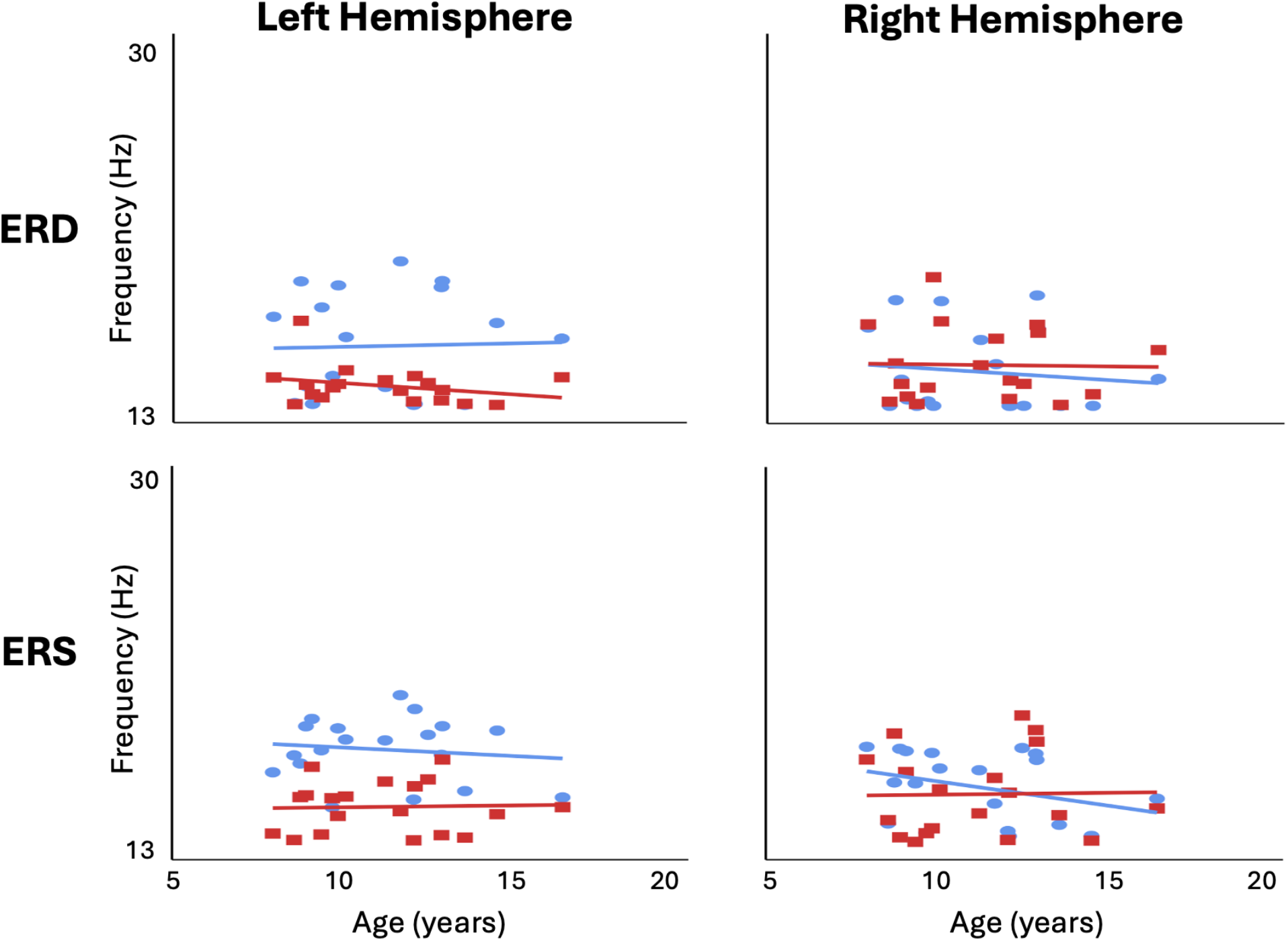
Peak frequencies for child group plotted by age. Blue ovals = manual task magenta squares = speech task. Linear regression lines are plotted separately for speech and manual data.

### Group differences

Group comparisons using nonparametric Kruskal-Wallis one-way ANOVA did not achieve statistical significance for either the manual or speech tasks: Manual (AD *Mdn* = 17.648 Hz, TD *Mdn* = 15.956 Hz, χ^2^(1,27) = 3.71, *p* = .054). Speech (AD *Mdn* = 15.653 Hz, TD *Mdn* = 14.897 Hz, χ^2^(1,27) = 1.42, *p* = .233).

## DISCUSSION

Much of the by now substantial literature on the beta band motor rhythm (for reviews see (28–32)) have considered it to reflect the operations of single mechanistic and functional entity. However there are good reasons to believe that the overall band is heterogenous and composed of distinct oscillations with different frequency signatures (29). This has been well-demonstrated for the basic subdivisions of high (> ∼20 Hz) versus low (< ∼20 Hz) beta, which have been reported to have distinct neural sources (33) (34) (35) and differential network involvement in motor preparation and execution (36). A few EEG (13, 37) and invasive electrophysiological (38) studies have reported evidence for yet finer grained beta frequency partitions associated with the movements of different bodily effectors. The EEG study of Neuper and Pfurtscheller (2001) estimated peak beta frequencies for manual (self-paced extension and flexion of the right index finger, measured at C3 electrode position overlying the hand representation) and foot (self-paced dorsiflexion of the right foot, measured at Cz overlying the foot representation) in healthy adults. These authors reported significantly greater peak frequency for foot movements (*M* = 21.5 Hz) compared to hand movements (M = 17.42 H, in close agreement with the present finger movement results, M = 17.88 Hz). Local field potential (LFP) evidence for a comparable beta effector specificity has also been reported in the human subthalamic nucleus, where lower limb movements elicited significantly greater activity in the higher beta range (24-31 Hz) than upper limb movements (38). Finally, Pfurtscheller et al. (37) reported limited EEG evidence (in a single participant) for a beta frequency distinction between different digits of the same hand. To our knowledge, these reports constitute the current literature on effector frequency specificity for the beta motor rhythm (although see (39) for electrocorticographic (ECoG) evidence of hand digit frequency specificity for gamma band oscillations that are phase-entrained by oscillations within the low (12-20 Hz) beta band).

The current results extend this evidence base to a speech-movement associated area situated in the middle portion of the precentral gyrus immediately ventral to the hand representation. Taken together, our results and those of Neuper and Pfurtscheller (2001) point to a spatial gradient of beta frequencies along the dorsal to ventral axis of this portion of the M1 motor strip, i.e. foot area ∼ 21.5 Hz, hand area ∼ 17.5 Hz, speech area ∼ 16.5 Hz. Viewed in the framework of dynamic systems, the existence of a spatial frequency gradient has important implications for oscillating networks, since a frequency difference between oscillators will limit the extent to which they can synchronize their oscillations, depending on the slope of the frequency gradient and the strength of coupling between oscillators. Models of neural network oscillations posit that network synchrony – defined as a stable phase relation between oscillators -- is a critical mechanism for neural communication and cooperation. On the other hand, the existence of a frequency difference (frequency detuning) between oscillators acts against synchronization by causing their phase relationship to change continuously (phase precession). These phenomena provide a conceptually powerful mechanistic solution for the overall problem of achieving differential control of bodily representations within a spatially continuous topographic map, by synchronizing neuronal activities within a representation, while segregating activities between representations by frequency detuning. At the present time these frequency-based mechanisms are strikingly absent from most discussions of M1 organization and function and merit renewed consideration in the current and ongoing reconceptualization of topographic motor representations.

The present results provide novel evidence for both hemispheric and developmental influences on beta oscillatory frequency mechanisms. Both children and adults showed clear task frequency differences in the left hemisphere, while in contrast to adults, children showed no significant task frequency differences in the right hemisphere. Following the logic of oscillatory frequency detuning described above, these results suggest an early developing mechanism of functional segregation in the left hemisphere in conjunction with a later developing mechanism in the right. Another intriguing group difference is the finding of significantly lower beta frequencies (for both tasks) in the right hemisphere of adults but not children, suggesting that the functional architecture of the mature brain is characterized by a more prominent separation of the activities of the two hemispheres.

### Conclusions

The present results provide novel evidence that the beta motor rhythm frequency is effector specific, showing significantly different peak frequencies within spatially contiguous regions of primary sensorimotor cortex associated with finger and speech movements, in both mature and developing human brains. These results bear on issues of considerable current interest within several distinct neuroscientific literatures, including ongoing enquiries into the mechanisms and functions of the beta rhythm itself; and recent investigations into the nature and organization of bodily representations within the primary motor cortex. Our data from children further suggest new insights into the development of motor control in the immature brain.

## DATA AVAILABILITY

Data are available upon request to the authors.

## AUTHOR CONTRIBUTIONS

AI, DC and BJ conceived and designed research, performed experiments, analyzed data, interpreted results of experiments, prepared figures, drafted manuscript, edited and revised manuscript, approved final version of manuscript.

## REFERENCES

1. Penfield W, Boldrey E. Somatic motor and sensory representation in the cerebral cortex of man as studied by electrical stimulation. Brain 60: 389–443, 1937. doi: 10.1093/brain/60.4.389.

2. Deo DR, Okorokova EV, Pritchard AL, Hahn NV, Card NS, Nason-Tomaszewski SR, Jude J, Hosman T, Choi EY, Qiu D, Meng Y, Wairagkar M, Nicolas C, Kamdar FB, Iacobacci C, Acosta A, Hochberg LR, Cash SS, Williams ZM, Rubin DB, Brandman DM, Stavisky SD, AuYong N, Pandarinath C, Downey JE, Bensmaia SJ, Henderson JM, Willett FR. A mosaic of whole-body representations in human motor cortex. bioRxiv: 2024.09.14.613041, 2024.

3. Gordon EM, Chauvin RJ, Van AN, Rajesh A, Nielsen A, Newbold DJ, Lynch CJ, Seider NA, Krimmel SR, Scheidter KM, Monk J, Miller RL, Metoki A, Montez DF, Zheng A, Elbau I, Madison T, Nishino T, Myers MJ, Kaplan S, Badke D’Andrea C, Demeter DV, Feigelis M, Ramirez JSB, Xu T, Barch DM, Smyser CD, Rogers CE, Zimmermann J, Botteron KN, Pruett JR, Willie JT, Brunner P, Shimony JS, Kay BP, Marek S, Norris SA, Gratton C, Sylvester CM, Power JD, Liston C, Greene DJ, Roland JL, Petersen SE, Raichle ME, Laumann TO, Fair DA, Dosenbach NUF. A somato-cognitive action network alternates with effector regions in motor cortex. Nature 617: 351–359, 2023. doi: 10.1038/s41586-023-05964-2.

4. Jensen MA, Huang H, Valencia GO, Klassen BT, Van Den Boom MA, Kaufmann TJ, Schalk G, Brunner P, Worrell GA, Hermes D, Miller KJ. A motor association area in the depths of the central sulcus. Nat Neurosci 26: 1165–1169, 2023. doi: 10.1038/s41593-023-01346-z.

5. Taheri A. The partial upward migration of the laryngeal motor cortex: A window to the human brain evolution. Brain Res 1834: 148892, 2024. doi: 10.1016/j.brainres.2024.148892.

6. Willett FR, Deo DR, Avansino DT, Rezaii P, Hochberg LR, Henderson JM, Shenoy KV. Hand knob area of premotor cortex represents the whole body in a compositional way. Cell 181: 1–14, 2020. doi: 10.1016/j.cell.2020.02.043.

7. Schieber MH. Constraints on somatotopic organization in the primary motor cortex. J Neurophysiol 86: 2125–2143, 2001. doi: 10.1152/jn.2001.86.5.2125.

8. Tremblay P, Brambati SM. A historical perspective on the neurobiology of speech and language: from the 19th century to the present. Front Psychol 15, 2024. doi: 10.3389/fpsyg.2024.1420133.

9. Thomas Yeo BT, Krienen FM, Sepulcre J, Sabuncu MR, Lashkari D, Hollinshead M, Roffman JL, Smoller JW, Zöllei L, Polimeni JR, Fischl B, Liu H, Buckner RL. The organization of the human cerebral cortex estimated by intrinsic functional connectivity. J Neurophysiol 106: 1125–1165, 2011. doi: 10.1152/jn.00338.2011.

10. Silva AB, Liu JR, Zhao L, Levy DF, Scott TL, Chang EF. A neurosurgical functional dissection of the middle precentral gyrus during speech production. J Neurosci 42: 8416–8426, 2022. doi: 10.1523/JNEUROSCI.1614-22.2022.

11. Lowet E, De Weerd P, Roberts MJ, Hadjipapas A. Tuning neural synchronization: The role of variable oscillation frequencies in neural circuits. Front Syst Neurosci 16: 908665, 2022. doi: 10.3389/fnsys.2022.908665.

12. Buzsaki G. Rhythms of the Brain. Oxford University Press, 2006.

13. Neuper C, Pfurtscheller G. Evidence for distinct beta resonance frequencies in human EEG related to specific sensorimotor cortical areas. Clin Neurophysiol 112: 2084–2097, 2001. doi: 10.1016/S1388-2457(01)00661-7.

14. Plate A, Hell F, Mehrkens JH, Koeglsperger T, Bovet A, Stanslaski S, Bötzel K. Peaks in the beta band of the human subthalamic nucleus: a case for low beta and high beta activity. J Neurosurg 136: 672–680, 2022. doi: 10.3171/2021.3.JNS204113.

15. Anastasopoulou I, Cheyne DO, Van Lieshout P, Johnson BW. Decoding kinematic information from beta-band motor rhythms of speech motor cortex: A methodological/analytic approach using concurrent speech movement tracking and magnetoencephalography. Front Hum Neurosci 18, 2024. doi: 10.3389/fnhum.2024.1305058.

16. Anastasopoulou I, Cheyne DO, Lieshout P van, Wilson PH, Ballard KJ, Johnson BW. A novel candidate neuromarker of central motor dysfunction in childhood apraxia of speech. bioRxiv: 2024.08.11.607491, 2024.

17. Veale JF. Edinburgh Handedness Inventory – Short Form: A revised version based on confirmatory factor analysis. Laterality Asymmetries Body Brain Cogn 19: 164–177, 2014. doi: 10.1080/1357650X.2013.783045.

18. Kado H, Higuchi M, Shimogawara M, Haruta Y, Adachi Y, Kawai J, Ogata H, Uehara G. Magnetoencephalogram systems developed at KIT. IEEE Trans Appl Supercond 9: 4057–4062, 1999. doi: 10.1109/77.783918.

19. Uehara G, Adachi Y, Kawai J, Shimogawara M, Higuchi M, Haruta Y, Ogata H, Kado H. Multi-channel SQUID systems for biomagnetic measurement. IEICE Trans Electron E86-C: 43–54, 2003.

20. Litvak V, Mattout J, Kiebel S, Phillips C, Henson R, Kilner J, Barnes G, Oostenveld R, Daunizeau J, Flandin G, Penny W, Friston K. EEG and MEG data analysis in SPM8. Comput Intell Neurosci 2011: 1–32, 2011. doi: 10.1155/2011/852961.

21. Frankford SA, Nieto-Castañón A, Tourville JA, Guenther FH. Reliability of single-subject neural activation patterns in speech production tasks. Brain Lang 212: 104881, 2021. doi: 10.1016/j.bandl.2020.104881.

22. Riecker A, Ackermann H, Wildgruber D, Meyer J, Dogil G, Haider H, Grodd W. Articulatory/phonetic sequencing at the level of the anterior perisylvian cortex: A functional magnetic resonance imaging (fMRI) study. Brain Lang 75: 259–276, 2000. doi: 10.1006/brln.2000.2356.

23. van Lieshout PHHM, Bose A, Square PA, Steele CM. Speech motor control in fluent and dysfluent speech production of an individual with apraxia of speech and Broca’s aphasia. Clin Linguist Phon 21: 159–188, 2007. doi: 10.1080/02699200600812331.

24. Jobst C, Ferrari P, Isabella S, Cheyne D. Brainwave: A matlab toolbox for beamformer source analysis of MEG data. Front Neurosci 12, 2018. doi: 10.3389/fnins.2018.00587.

25. Yousry TA, Schmid UD, Alkadhi H, Schmidt D, Peraud A, Buettner A, Winkler P. Localization of the motor hand area to a knob on the precentral gyrus. A new landmark. Brain 120: 141–157, 1997. doi: 10.1093/brain/120.1.141.

26. Bickel PJ, Sakov A. On the choice of m in the m out of n bootstrap and confidence bounds for extrema. Stat Sin 18: 967–85, 2008.

27. Muthukumaraswamy SD, Singh KD. Visual gamma oscillations: The effects of stimulus type, visual field coverage and stimulus motion on MEG and EEG recordings. NeuroImage 69: 223–230, 2013. doi: 10.1016/j.neuroimage.2012.12.038.

28. Barone J, Rossiter HE. Understanding the role of sensorimotor beta oscillations. Front Syst Neurosci 15, 2021. doi: 10.3389/fnsys.2021.655886.

29. Schmidt R, Herrojo Ruiz M, Kilavik BE, Lundqvist M, Starr PA, Aron AR. Beta oscillations in working memory, executive control of movement and thought, and sensorimotor function. J Neurosci 39: 8231–8238, 2019. doi: 10.1523/JNEUROSCI.1163-19.2019.

30. Cheyne DO. MEG studies of sensorimotor rhythms: a review. Exp Neurol 245: 27–39, 2013. doi: 10.1016/j.expneurol.2012.08.030.

31. Kilavik BE, Zaepffel M, Brovelli A, MacKay WA, Riehle A. The ups and downs of β oscillations in sensorimotor cortex. Exp Neurol 245: 15–26, 2013. doi: 10.1016/j.expneurol.2012.09.014.

32. Lundqvist M, Miller EK, Nordmark J, Liljefors J, Herman P. Beta: Bursts of cognition. Trends Cogn Sci 28: 662–676, 2024. doi: 10.1016/j.tics.2024.03.010.

33. Nougaret S, López-Galdo L, Caytan E, Poitreau J, Barthelemy FV, Kilavik BE. Distinct sources and behavioral correlates of macaque motor cortical low and high beta. bioRxiv: 2023.03.28.534535, 2023.

34. West TO, Berthouze L, Halliday DM, Litvak V, Sharott A, Magill PJ, Farmer SF. Propagation of beta/gamma rhythms in the cortico-basal ganglia circuits of the parkinsonian rat. J Neurophysiol 119: 1608–1628, 2018. doi: 10.1152/jn.00629.2017.

35. López-Azcárate J, Tainta M, Rodríguez-Oroz MC, Valencia M, González R, Guridi J, Iriarte J, Obeso JA, Artieda J, Alegre M. Coupling between beta and high-frequency activity in the human subthalamic nucleus may be a pathophysiological mechanism in parkinson’s disease. J Neurosci 30: 6667–6677, 2010. doi: 10.1523/JNEUROSCI.5459-09.2010.

36. Chandrasekaran C, Bray IE, Shenoy KV. Frequency shifts and depth dependence of premotor beta band activity during perceptual decision-making. J Neurosci 39: 1420–1435, 2019. doi: 10.1523/JNEUROSCI.1066-18.2018.

37. Pfurtscheller G, Woertz M, Krausz G, Neuper C. Distinction of different fingers by the frequency of stimulus induced beta oscillations in the human EEG. Neurosci Lett 307: 49–52, 2001. doi: 10.1016/S0304-3940(01)01924-3.

38. Tinkhauser G, Shah SA, Fischer P, Peterman K, Debove I, Nygyuen K, Nowacki A, Torrecillos F, Khawaldeh S, Tan H, Pogosyan A, Schuepbach M, Pollo C, Brown P. Electrophysiological differences between upper and lower limb movements in the human subthalamic nucleus. Clin Neurophysiol 130: 727–738, 2019. doi: 10.1016/j.clinph.2019.02.011.

39. Miller KJ, Zanos S, Fetz EE, Nijs M den, Ojemann JG. Decoupling the cortical power spectrum reveals real-time representation of individual finger movements in humans. J Neurosci 29: 3132–3137, 2009. doi: 10.1523/JNEUROSCI.5506-08.2009.

